# A Binary Matrix Method to Enumerate, Hierarchically Order and Structurally Classify Peptide Aggregation

**DOI:** 10.1101/2021.11.29.470297

**Authors:** Amol Tagad, Reman Kumar Singh, G. Naresh Patwari

## Abstract

Protein aggregation is a common and complex phenomenon in biological processes, yet a robust analysis of this aggregation process remains elusive. The commonly used methods such as center-of-mass to center-of-mass (*COM*–*COM*) distance, the radius of gyration (*R*_*g*_), hydrogen bonding (HB) and solvent accessible surface area (SASA) do not quantify the aggregation accurately. Herein, a new and robust method that uses an aggregation matrix (AM) approach to investigate peptide aggregation in a MD simulation trajectory is presented. A *nxn* two-dimensional aggregation matrix (AM) is created by using the inter-peptide *C*_*α*_–*C*_*α*_ cut-off distances which are binarily encoded (0 or 1). These aggregation matrices are analyzed to enumerate, hierarchically order and structurally classify the aggregates. Comparison of the present AM method suggests that it is superior to the HB method since it can incorporate non-specific interactions and *R*_*g*_, *COM*–*COM* methods since the cut-off distance is independent of the length of the peptide. More importantly, the present method can structurally classify the peptide aggregates, which the conventional *R*_*g*_, *COM*– *COM* and HB methods fail. The unique selling point of this method is its ability to structurally classify peptide aggregates using two-dimensional matrices.

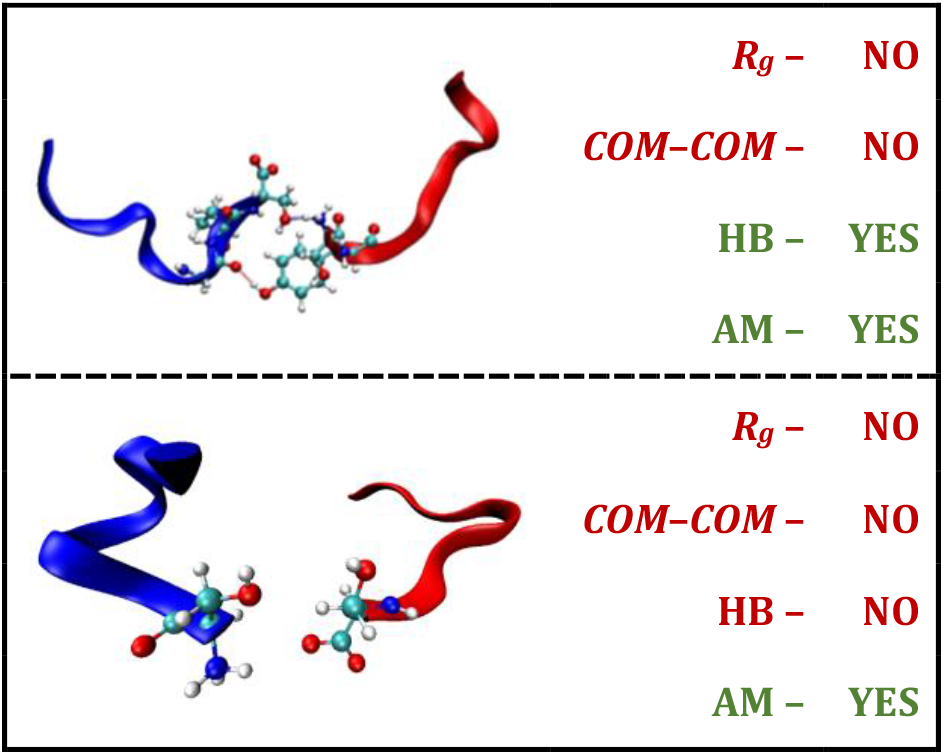

## INTRODUCTION

Protein-protein (or inter-protein) interactions are important in many biological processes,^1–3^ such as translation,^4^ transcription factors,^5^ cell signalling,^6–8^ protein synthesis,^9,10^ and many others. These inter-protein interactions are primarily driven by high hydrophobicity, low charge density, and high β-sheet propensity.^11,12^ Many interprotein interactions involve folded proteins, however, in some cases, these interactions are formed by disordered or structureless proteins and peptides, which aggregate to form fibrillar structures and result in amyloid fibrils.^13–17^ In amyloid fibrils are insoluble aggregates formed as cross β-sheet patterns, where the β-sheet backbone is perpendicular to the fibril axis^8,18–20^ These protein aggregates are associated with diseases like Alzheimer’s,^21–23^ Parkinson’s,^16,24,25^ Huntington’s, type II diabetes mellitus,^7,24,25^ and several others, therefore, the atomic level understanding of the aggregation process is vitally important. Apart from several experimental methodologies,^2,16,26–28^ molecular dynamics (MD) simulations have been frequently used to investigate peptide and protein aggregation to understand the aggregation pathways along with their structural characterization and dynamic behavior. In this regard, several physical and chemical parameters such as center-of-mass to center-of-mass (*COM*–*COM*) distance, the radius of gyration (*R*_*g*_), the dihedral angle between the peptide chains, hydrogen bonding (HB), solvent accessible surface area (SASA), and contact matrix have been primarily used to probe the aggregation in a MD trajectory.^22,29,30^ While the above mentioned methods have been utilized to investigate peptide/protein aggregation, the global methods such as *COM*–*COM, R*_*g*_ and dihedral angle between the peptide chains do not consider the residue level interaction. On the other hand, the residue level methods such as HB, SASA and contact matrix are in general qualitative and not quantitative. More importantly, none of these methods, except the HB method, provide information on the hierarchy of aggregation, which can be utilized to estimate the propensity to form compact and energetically favorable aggregates. However, none of the conventional methods including HB can differentiate the structurally ordering, such as the formation of parallel and antiparallel aggregates. Herein, a new and robust method, which uses an aggregation matrix approach to investigate peptide aggregation in a MD simulation trajectory is proposed. This method utilizes the inter-peptide *C*_*α*_–*C*_*α*_ cut-off distances which are binarily encoded (0 or 1) to create *n*x*n* two-dimensional matrices, which are further analyzed to enumerate, hierarchically order and structurally classify the aggregates. The unique feature of the proposed method is its ability to structurally classify peptide aggregates, which cannot be carried out using existing methods. In the present work, the aggregation propensity of six known low complexity domain (LCD) peptides,^11,19,39,40,31–38^ using the aggregation matrix approach is investigated and compared with the other methods.

## METHODOLOGY

In this work, six LCD peptides with the sequences NKGAII (3Q2X),^41^ STGGYG (6BZP),^2^ SYSGYS (6BWZ),^2^ SYSSYGQS (6BXV),^2^ GYNGFG (6BXX),^2^ and GFGNFGTS (6BZM)^2^ were considered. The initial conformation was taken directly from the PDB and the dimers, trimers and tetramers were generated by replicating the monomers and placing them such that the surfaces of the monomers is at a distance of 10Å in a cubic box with the dimension such that the peptides do not interact with their periodic image during the simulation. The peptides were solvated by TIP3P water model^42^ and Na^+^ and Cl^−^ ions were added to neutralize the system. All the simulations were carried out with CHARMM36 force field^43^ using GROMACS 2019 software package.^44^ The potential energy of the system was minimized for 5000 steps using the steepest descent method.^45^ During the minimization a 10 Å cut-off was used for van der Waals and electrostatic interactions. The particle mesh Ewald (PME) method was used to model the electrostatic interactions.^46^Additionally, the bonds were constrained using the LINCS algorithm.^47^ Following energy minimization, each system was equilibrated in NVT and NPT ensembles for 200 ps and 1000 ps, respectively.^48^ The velocity rescaling thermostat^49^ and Parrinello-Rahman barostat^50^ were used to maintain 300 K temperature and 1 bar pressure with a coupling constant of 0.4 ps. Finally, the production simulation using NPT condition at 300 K and 1 bar pressure was carried and the equations of motion were integrated using the leapfrog algorithm with a time step of 2 fs. The simulation time was 2 μs for each of the systems containing two, three and four peptide units, and the coordinates were saved every 10 ps during the simulation and the data was further analysed.

The formation of the peptide aggregates was analyzed using inter-peptide pair-wise *C*_*α*_–*C*_*α*_ distances. The inter-peptide *C*_*α*_–*C*_*α*_ distances were cast as binary logic of 1 or 0 with a cut-off distance criterion and an aggregation matrix was generated. The entries in the aggregation matrix would be 1 or 0 depending on whether the inter-peptide *C*_*α*_–*C*_*α*_ distance was less than or equal or greater to the cut-off distance, respectively. For a *n*-mer peptide the aggregation matrix would be of the order *n*x*n* with the matrix elements being 0 or 1. This method is similar to the adjacency matrix for contact formation, but the advantage of using an aggregation matrix is that each element *a*_*ij*_ binarily encodes the presence/absence of inter-peptide interaction between the *C*_*α*_ of *i*_th_ residue of the first (A) peptide to the *C*_*α*_ of the *j*_th_ residue of the second (B) peptide. The aggregation matrices were generated for each snapshot over the entire trajectory. This method scales as *n*^2^ for the dimer, where *n* is the number of residues in the peptide. For a 2 μs trajectory (200,000 snapshots) the time taken to calculate the aggregation matrix and the corresponding aggregation number for a dimer of 8-mer peptide is about 6 minutes using Intel-Xeon 8 core E3 processor, which makes it computationally very tractable.

The formation of an aggregated dimer can be defined when the sum of all the elements of the aggregation matrix, the aggregation number, which is defined as

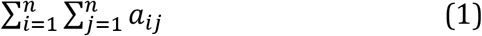

is non-zero. The aggregation number can also be used to define the hierarchy of aggregation, wherein a larger aggregation number indicates a more compact structure. Further, the formation of a specific type of aggregate can also be inferred from the aggregation matrix. For instance, the formation of a quasi-parallel dimer configuration can be recognized by the sum of the diagonal elements, which is defined as

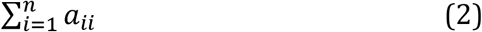

On the other hand, a quasi-anti-parallel dimer configuration can be recognized by the sum of the anti-diagonal elements, which is defined as

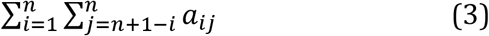

In the present analysis, the formation of parallel/anti-parallel configurations is evaluated with the constraint of the sum of the diagonal/anti-diagonal elements being greater than or equal to 3. The choice of minimum value of 3 for such classification is based on the fact that a lower value would not be able to distinguish between the parallel/anti-parallel structures at marginally higher *C*_*α*_–*C*_*α*_ cut-off distances and a smaller *C*_*α*_–*C*_*α*_ cut-off distances will artificially lower the aggregation propensity. Similarly, several shifted parallel and shifted anti-parallel dimer configurations can be defined by either increasing or decreasing the index *j* relative to index *i*, such as

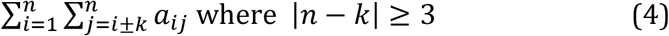

for shifted parallel, and

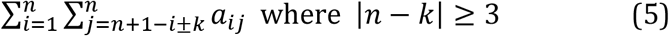

for shifted anti-parallel configurations. Other configurations such as T-shaped and X-shaped can also be identified from the *n*x*n* aggregation matrix. For the T-shaped dimer, only one element of either first row (*a*_1*j*,_) or first column (*a*_*i*1_) or last row (*a*_*nj*_) or last column (*a*_*in*_) must be 1 and all the other elements must be 0s. On the other hand, for a X-shaped structure, the elements of a *m*x*m* (where *m* ≪*n*) block must be 1s and all the other elements must be 0s. A trimer has three identical peptides, however, for representation purposes, three of the peptides are indexed as A, B, and C. In a system with three peptides aggregation leads to the formation of dimers and trimers. In this case, the formation of three independent dimers (AB, BC and CA) are analyzed using the aggregation number given in equation (1). Further, the formation of a trimer is recognized by snapshots/frames that contain two or more independent dimers. The hierarchy in the trimers can be defined by the sum of elements in all the three independent dimer aggregation matrices. This methodology was extended to tetramers as well, in which six independent dimers (AB, AC, AD, BC, BD, and CD), four independent trimers (ABC, ABD, ACD, and BCD) and a tetramer are formed. A trimer is formed by snapshots/frames that contain two or more independent dimers with one common peptide unit (five out of six combinations will satisfy this condition). Finally, the formation of tetramers was based on the inter-peptide *C*_*α*_–*C*_*α*_ distance being less than the cut-off value across the three sets of mutually exclusive dimers (AB-CD, AC-BD, and AD-BC) in a single snapshot. Further, the propensity to form ordered structures such as parallel, shifted parallel, anti-parallel and shifted anti-parallel, for trimers and tetramers were analyzed based on equations (2-5).

## RESULTS

The aggregation behavior of the six LCD peptides considered in the present work was analyzed based on the aggregation matrix method, which was generated throughout the trajectory with inter-peptide *C*_*α*_–*C*_*α*_ cut-off distances of 0.50, 0.55, 0.60 0.65, 0.70 and 0.75 nm. Figure 1 shows the dimer population as a function of *C*_*α*_–*C*_*α*_ cut-off distance for all the peptides in a system containing two peptide units and the data is listed in Tables 1 and S1. The dimer population changes significantly in the cut-off distance range of 0.5 to 0.6 nm range, thereafter the population change is gradual. The convergence for the dimerization was checked over the entire trajectory in cumulative segments of 400 ns with *C*_*α*_–*C*_*α*_ cut-off distance of 0.55 nm (Table S2, see the SI). It can be noticed from Figure 1A that the extent of dimer population depends on the amino acid sequence, with the peptides SYSSYGQS and STGGYG having the highest and lowest propensity to form dimers, respectively. In general, the propensity to aggregate is higher for the 8-mers compared to 6-mers. The formation of dimers was also analyzed in terms of inter-peptide hydrogen bonds, shown in Figure 1B, and the results indicate very similar trends for various peptide sequences relative to the aggregation matrix method except for SYSGYS peptide. Further, the cumulative population of all the hydrogen-bonded dimers is similar to the dimer population obtained using *C*_*α*_–*C*_*α*_ cut-off distance of 0.55nm (Table S3, see the SI), which indicates that the aggregation is primarily hydrogen bond driven.

**Table 1:**
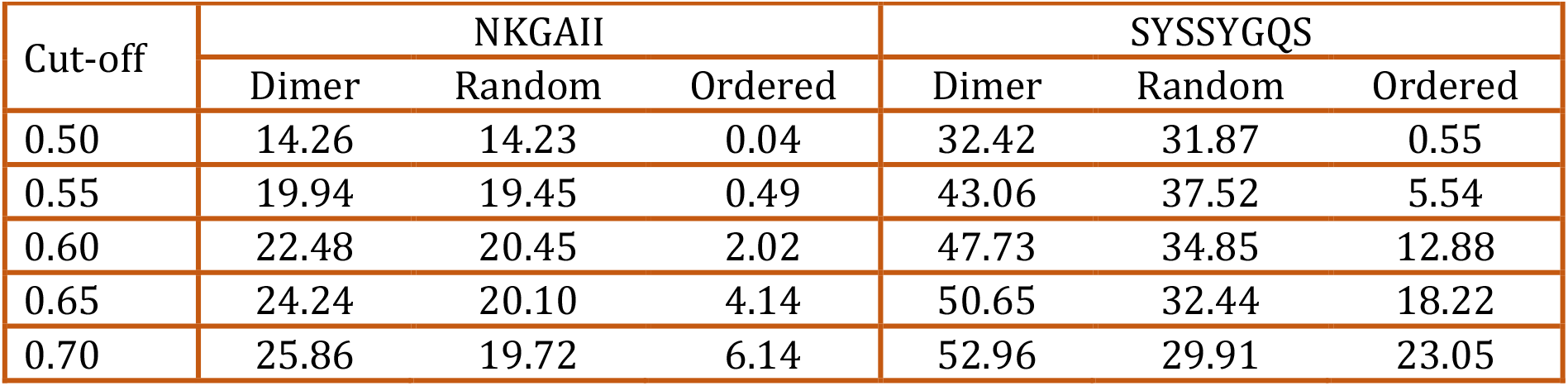
Representative dimer population (%) along with the population of ordered and random configurations at various inter-peptide *C*_*α*_–*C*_*α*_ cut-off distances (nm).

**Figure 1.**
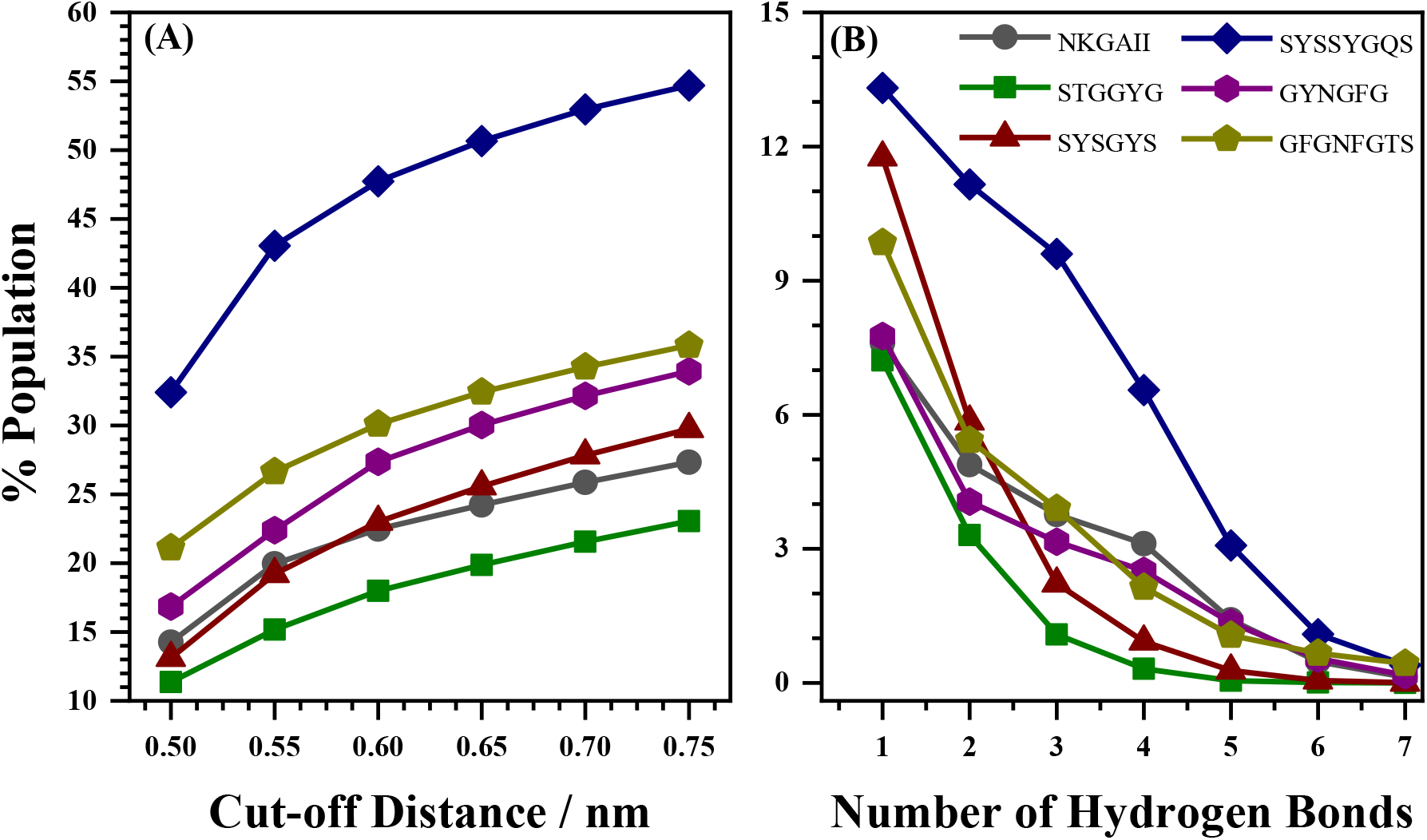
Dimer population (A) calculated using aggregation matrix as a function of *C*_*α*_– *C*_*α*_ cut-off distance and (B) as a function of the number of inter-peptide hydrogen bonds. Note that the legends in both the plots are the same.

One of the important consequences of using the aggregation matrix method is its ability to hierarchically order the structures based on the sum of all the entries in the aggregation matrix, the aggregation number. A larger aggregation number indicates the formation of a more compact structure. Figure S1 (see the SI) depicts the plots of dimer population as a function of aggregation number, which shows the presence of bimodal distribution for three peptide sequences (NKGAII, SYSSYGQS and GFGNFGTS) and roughly monotonic distribution for the other three peptides (STGGYG, SYSGYS and GYNGFG). The appearance of bimodal distribution in some cases can possibly be attributed to a lower probability of some contacts due to steric clashes of side chains. Significantly, the propensity to form a compact dimer is maximum for the SYSSYGQS peptide, which is in line with the propensity of this peptide to dimerize. The principal utility of the aggregation matrices is their ability to structurally classify peptide aggregates, which is elaborated in the methodology section and illustrated in Figure 2. In the present scenario, the ordering of peptides aggregates is based on the formation of parallel (and shifted parallel) or anti-parallel (and shifted anti-parallel) dimers, which are common structural motifs in various β-sheet secondary structures of proteins. Figure 2 also illustrates the classification of various structural configurations based on the appearance of the aggregation matrix. On the contrary, the hydrogen bond matrices do not give any discernible pattern for structure classification (Table S4, see the SI). In the present analysis, all the parallel, anti-parallel, shifted parallel and shifted anti-parallel dimers are pooled as ordered aggregates while the remaining population can be considered as random (non-specific) aggregates. In general, the propensity to form random aggregates is higher than the ordered aggregates, however, within the ordered aggregates, the propensity to form anti-parallel / shifted anti-parallel configurations is higher than parallel / shifted parallel configurations. This observation is in excellent agreement with the formation of antiparallel beta-sheets in peptide aggregates.^2,39,51^

**Figure 2.**
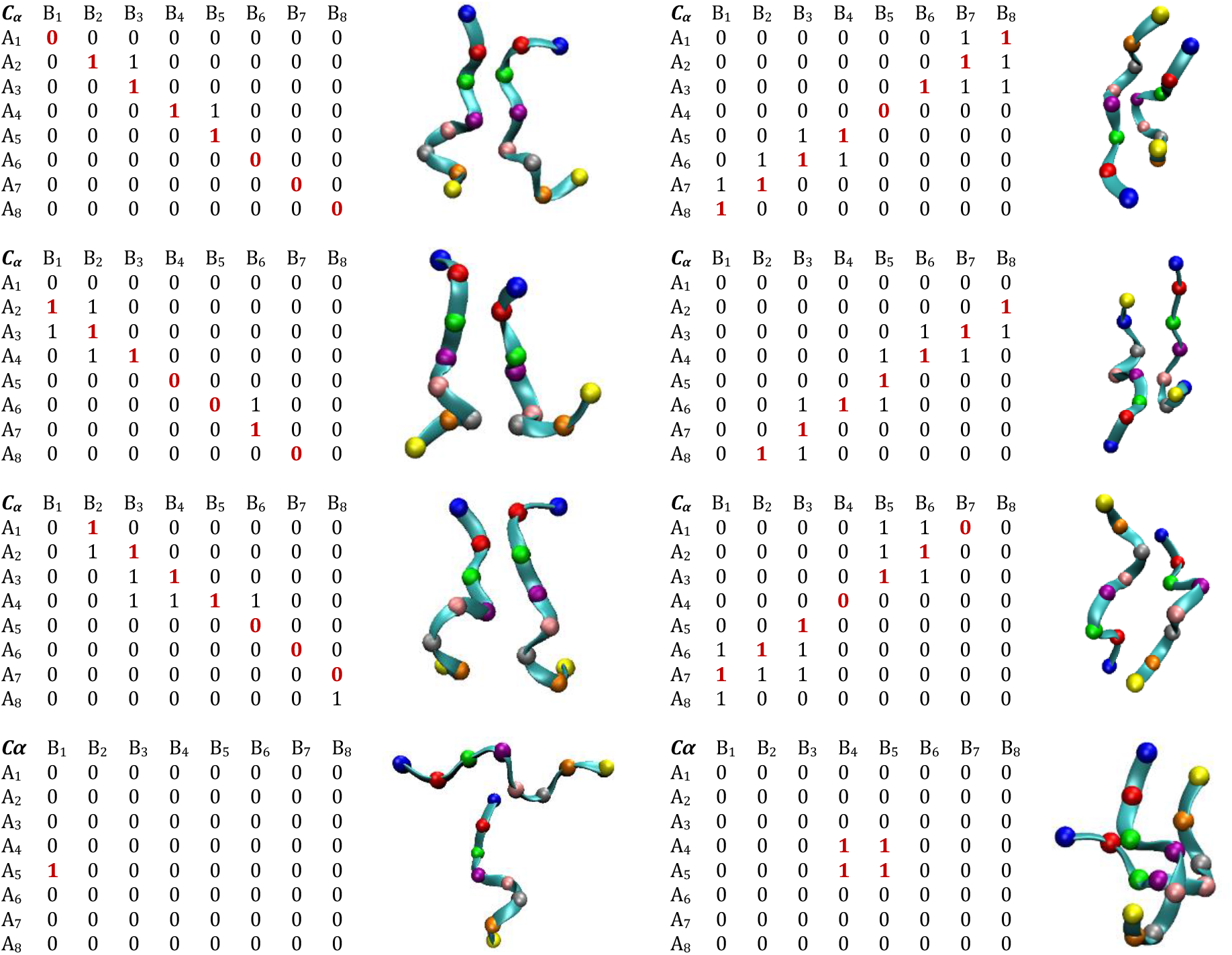
The matrix representations and the corresponding structures for the parallel/shifted parallel and T-shaped (left panel); anti-parallel / shifted anti-parallel and X-shaped (right panel) configurations of the SYSSYGQS dimer with the *C*_*α*_–*C*_*α*_ cutoff distance of 0.6 nm.

The role of side-chains (SC) in the dimerization of the SYSSYGQS peptide was evaluated by calculating the SC aggregation matrix, which is formed by the binary indexing of the distance between inter-peptide heavy atoms with a cut-off distance of 0.4 nm._52_ Figure 3 depicts the comparison of the data obtained from *C*_*α*_ (0.55 nm) and SC (0.4 nm) aggregation matrices. The green bars indicate the frequency of aggregation calculated over the snapshots that are common to *C*_*α*_ and SC aggregation matrices, while the red bars indicate the frequency of aggregation over the snapshots that are present in SC aggregation matrices but absent in *C*_*α*_ aggregation matrices. The black bar indicates the frequency of aggregation with no SC interactions. The data presented in Figure 4 illustrates that a large fraction (73.9 %) of aggregation is backbone driven and includes interaction between the side chains, while a smaller fraction (23.0 %) of aggregation is only due to side-chain interactions. On the other hand, a very small fraction (3.1%) of the dimer population due to interaction of the backbone and completely excludes inter-peptide side-chain interactions. Alternatively, the dimer population calculated using SC aggregation matrix (about 54.1%) can be recovered by increasing the *C*_*α*_–*C*_*α*_ cut-off distance to 0.75 nm (dimer population of 54.7%), which suggests that the dimer structures with the interaction between the side-chains can be assimilated by increasing the *C*_*α*_–*C*_*α*_ cut-off distance. However, with the increase in the *C*_*α*_–*C*_*α*_ cut-off distance, the aggregation matrix is not suitable for structural classification and structures of such side-chain based aggregates cannot be unambiguously characterized.

**Figure 3.**
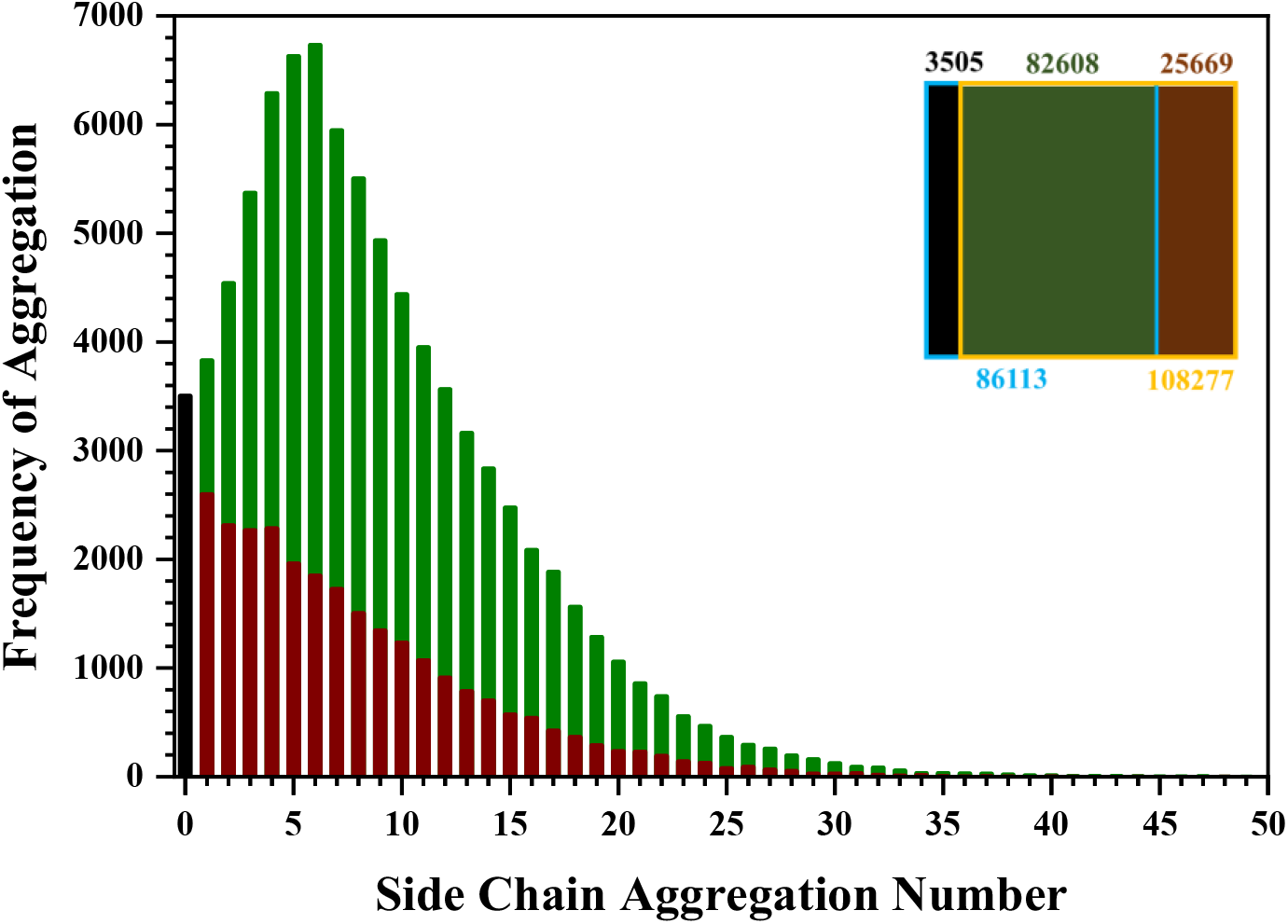
Histogram showing the frequency of aggregation for the inter-peptide sidechain (SC) aggregation number for the heavy atoms with a cut-off distance of 0.4 nm. The green bars indicate the frequency of aggregation over the snapshots that are common to *C*_*α*_ (0.55 nm) and SC aggregation matrices, while the brown bars indicate frequency aggregation over the snapshots that are present in SC aggregation matrices but absent in *C*_*α*_ (0.55 nm) aggregation matrices. The black bar indicates the frequency of aggregation with no SC interactions. The inset shows an overlap diagram with blue and yellow boxes representing snapshots with non-zero *C*_*α*_ and SC aggregation matrices. The green, brown and black regions represent the sum of corresponding cumulative frequency in the histogram. The colour-coded numbers represent the number of snapshots in each case out of a total of 200,000 snapshots. This overlap diagram clearly brings out the dominant contribution of the backbone to the aggregation.

**Figure 4.**
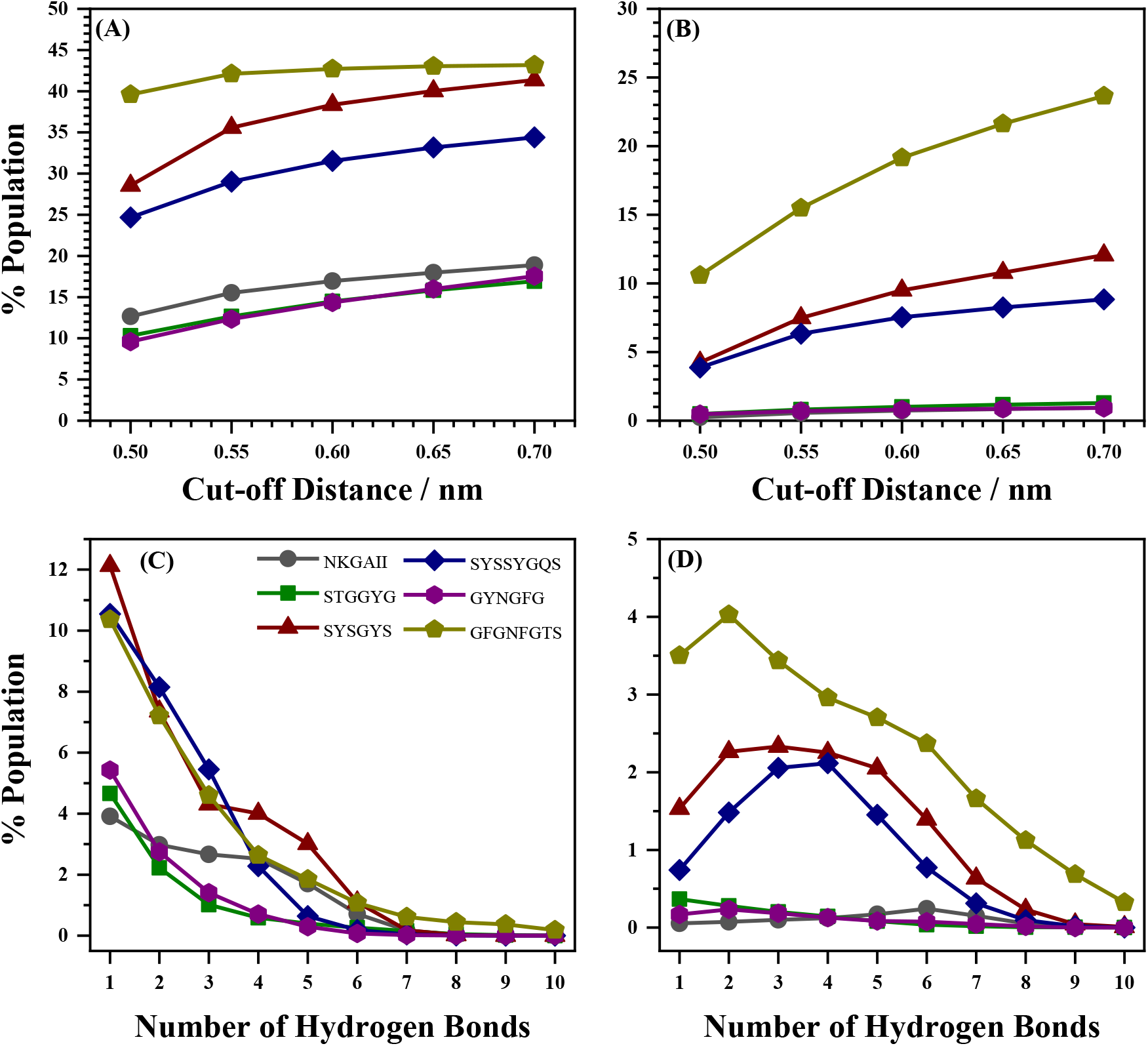
The population of (A) dimer and (B) trimers as a function of *C*_*α*_–*C*_*α*_ cut-off distance in a system containing three peptides units. In (A) the dimer population is the cumulative sum of all the three AB, BC and AC dimers while removing common frames. In (B) the trimer population was evaluated by only considering the common frames for at least two dimers. The population of (C) dimers and (D) trimers as a function of number of inter-peptide hydrogen bonds. The population of dimers and trimers are mutually exclusive. Note that the legends in all the plots are same.

In a system consisting of three peptides, the trimerization was analyzed by decomposing the trajectory into three pairs of dimers and the formation of the trimer was deciphered based on snapshots/frames with at least two sets of dimers. The population of the dimers and the trimers are mutually exclusive and are shown in Figure 4 as a function of *C*_*α*_–*C*_*α*_ cut-off distance and listed in Table S5 (see the SI). Interestingly, in the system containing three peptide units, the propensity to form either dimers or trimers showed a considerably different trend than the system containing two peptides. In this case, the sequence dependence on the propensity for trimer formation was GFGNFGTS > SYSGYS > SYSSYGQS and for the other three peptides, the timer population is negligible. Further, the aggregation behavior of trimers was analyzed in terms of inter-peptide hydrogen bonds, also shown in Figure 4, and the results indicate very similar trends relative to the aggregation matrix method, which once again suggests that the peptide aggregation is primarily hydrogen bond driven. Finally, in a system containing four peptide units, the formation of dimers, trimers and tetramers were evaluated and the results are shown in Figure 5 as a function of *C*_*α*_–*C*_*α*_ cut-off distance and listed in Table S6 (see the SI). The hierarchical ordering of aggregation (formation of dimers and trimers in a system containing three peptide units and formation of dimers, trimers and tetramers in a system containing four peptide units) has the same sequence dependence (Figure S1, see the SI) and is comparable to the hydrogen bonding propensity. Even in the case of trimers and tetramers, the appearance of bimodal distribution of number of *C*_*α*_–*C*_*α*_ contacts, in some cases, can possibly be attributed to a lower probability of some contacts due to steric clashes of side chains. A two-dimensional aggregation matrix approach to analyze the peptide aggregation in a MD trajectory for systems consisting of up to four peptide units is a useful and convenient methodology. As noted earlier, this method is capable of enumerating, hierarchically ordering and structurally classifying the aggregates which can be illustrated in Figure 6, wherein the formation of tetramer can be mediated by sequential addition of a monomer to form dimer, trimer and tetramer. Alternatively, the formation of a tetramer can also be formed by the aggregation of two dimers.

**Figure 5.**
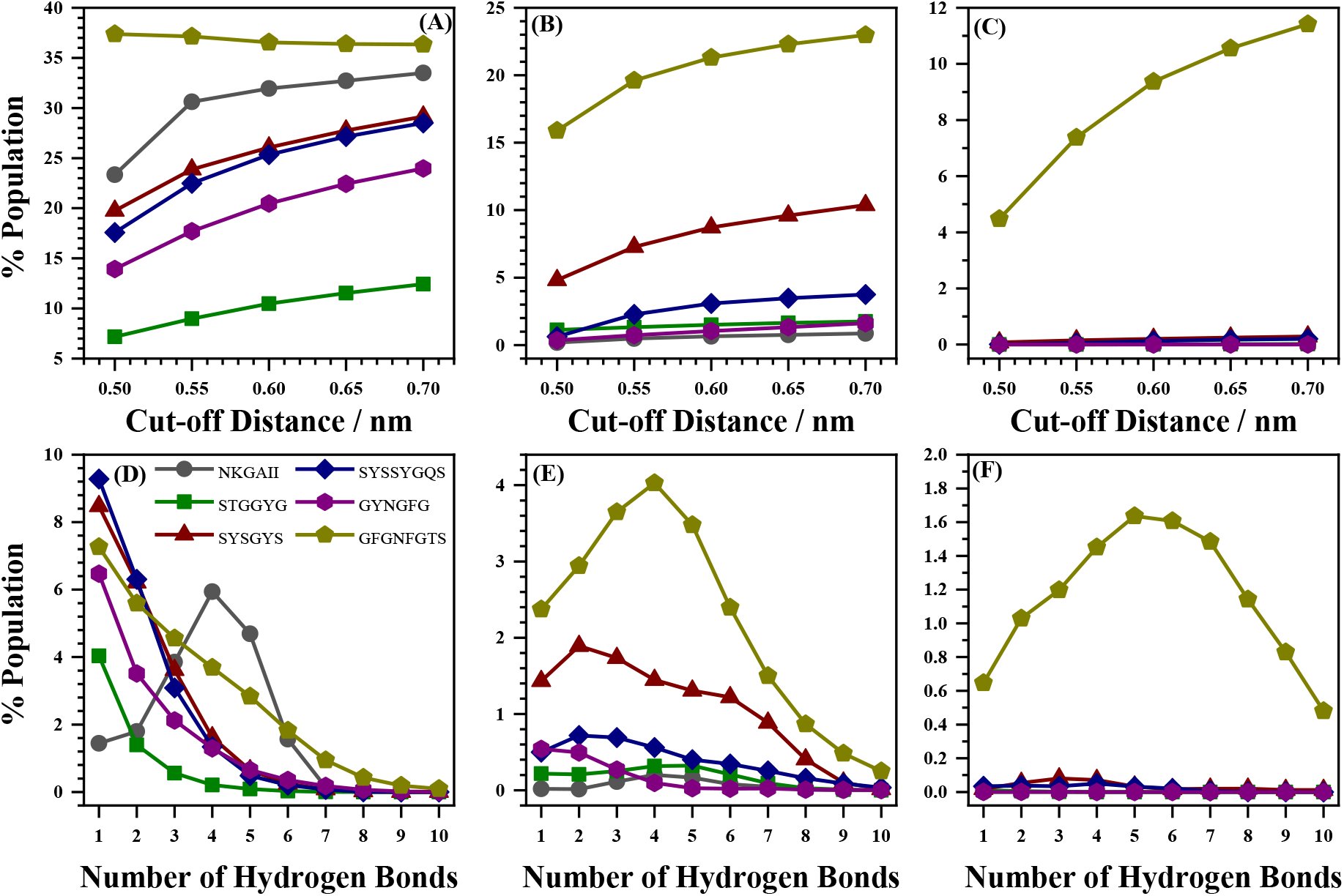
The population of (A) dimer, (B) trimers and (C) tetramers as a function of *C*_*α*_–*C*_*α*_ cut-off distance in a system containing four peptides units. In (A) the dimer population is the cumulative sum of all the six dimers AB, AC, AD, BC, BD and CD dimers while removing common frames. In (B) the trimer population was evaluated by only considering the common frames for at least two dimers with a common peptide unit (five out of six combinations will satisfy this condition), In (C) the tetramer formation evaluated based on the inter-peptide *C*_*α*_–*C*_*α*_ distance being less than the cut-off value across the three sets of mutually exclusive dimers (AB-CD, AC-BD, and AD-BC) in a snapshot. Population of (D) dimer, (E) trimers and (F) tetramers as a function of number of inter-peptide hydrogen bonds. The population of dimers, trimers and tetramers are mutually exclusive. Note that the legends in all the plots are the same.

**Figure 6.**
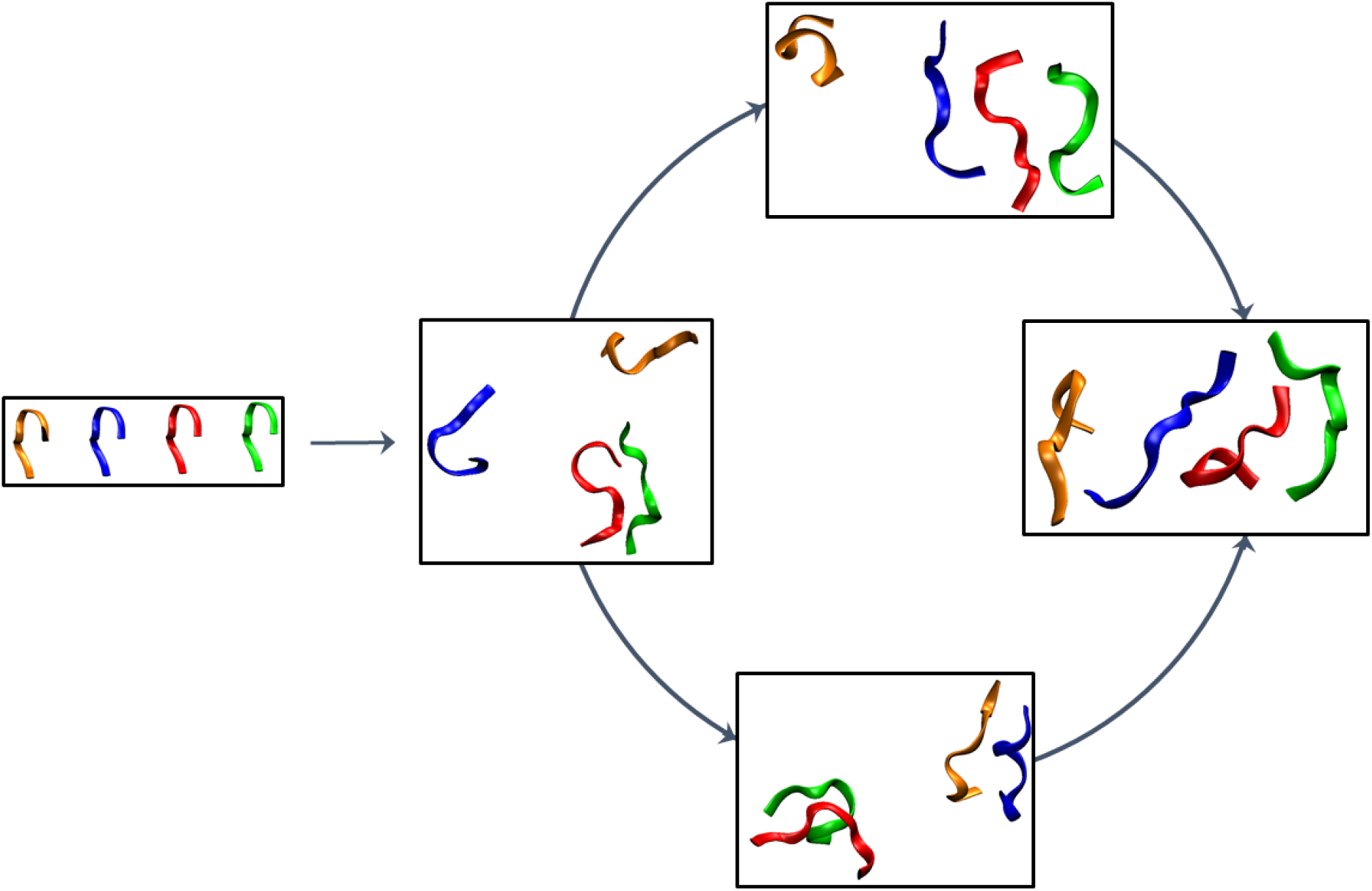
Snapshots of progression of aggregation in a system containing four peptide units of GFGNFGTS. Each peptide unit is colour coded for easy visualization. The formation of tetramer follows two pathways, first, step-wise addition, and second, dimerization of dimer.

## DISCUSSION

Commonly used methods to characterize peptide aggregation in the literature use radius of gyration (*R*_*g*_) and center-of-mass to center-of-mass distance (*COM*–*COM*).^22,29,30^ Figure 7 shows the comparison of the present aggregation matrix method with the *R*_*g*_ and *COM*– *COM* methods. In general, the *COM*–*COM* method grossly underestimates the propensity of aggregation with reasonable cut-off distances. On the other hand, the *R*_*g*_ method shows a comparable propensity of aggregation at a cut-off distance of 0.7 nm for the 6-mer peptides and substantially underestimates for 8-mer at the same cut-off distance of 0.7 nm. Whilst the *R*_*g*_ method can be used to estimate the propensity of aggregation, the cut-off distance varies based on the length of the peptide and cannot be uniformly applied across all peptide sequences. Further, Figure 8 illustrates the formation of two different end-to-end dimers for the SYSSYGQS peptide with very large values of *R*_*g*_ and *COM–COM*. These two aggregates, in general, are likely to be overlooked by both the *R*_*g*_ and *COM– COM* methods. On the other hand, these two aggregates are enumerated by the aggregation matrix method even if the structure is devoid of any hydrogen bonds. It is, therefore, essential to realize that *R*_*g*_ and *COM*–*COM* methods sample the global properties of the aggregate, while the aggregation matrix and hydrogen bonding methods sample the aggregation behavior at the residue level. Moreover, the aggregation matrix method is also capable of capturing non-specific (read non-hydrogen-bonded) dimers. The unique selling point (USP) of the aggregation matrix method is its ability to structurally classify peptide aggregation. The redundancy in the populations of ordered aggregates following structural classification increases with an increase in the *C*_*α*_–*C*_*α*_ cut-off distance as can be seen from Table S1, therefore caution must be exercised while interpreting the data corresponding to ordered aggregates. In view of the observed statistics and data redundancy in the ordered structures, a *C*_*α*_–*C*_*α*_ cut-off distance of around 0.55 nm is recommended, which is in good agreement with the *C*_*α*_–*C*_*α*_ distances observed in a Aβ(1–42) amyloid fibril (PDB 2NAO).^53^

**Figure 7.**
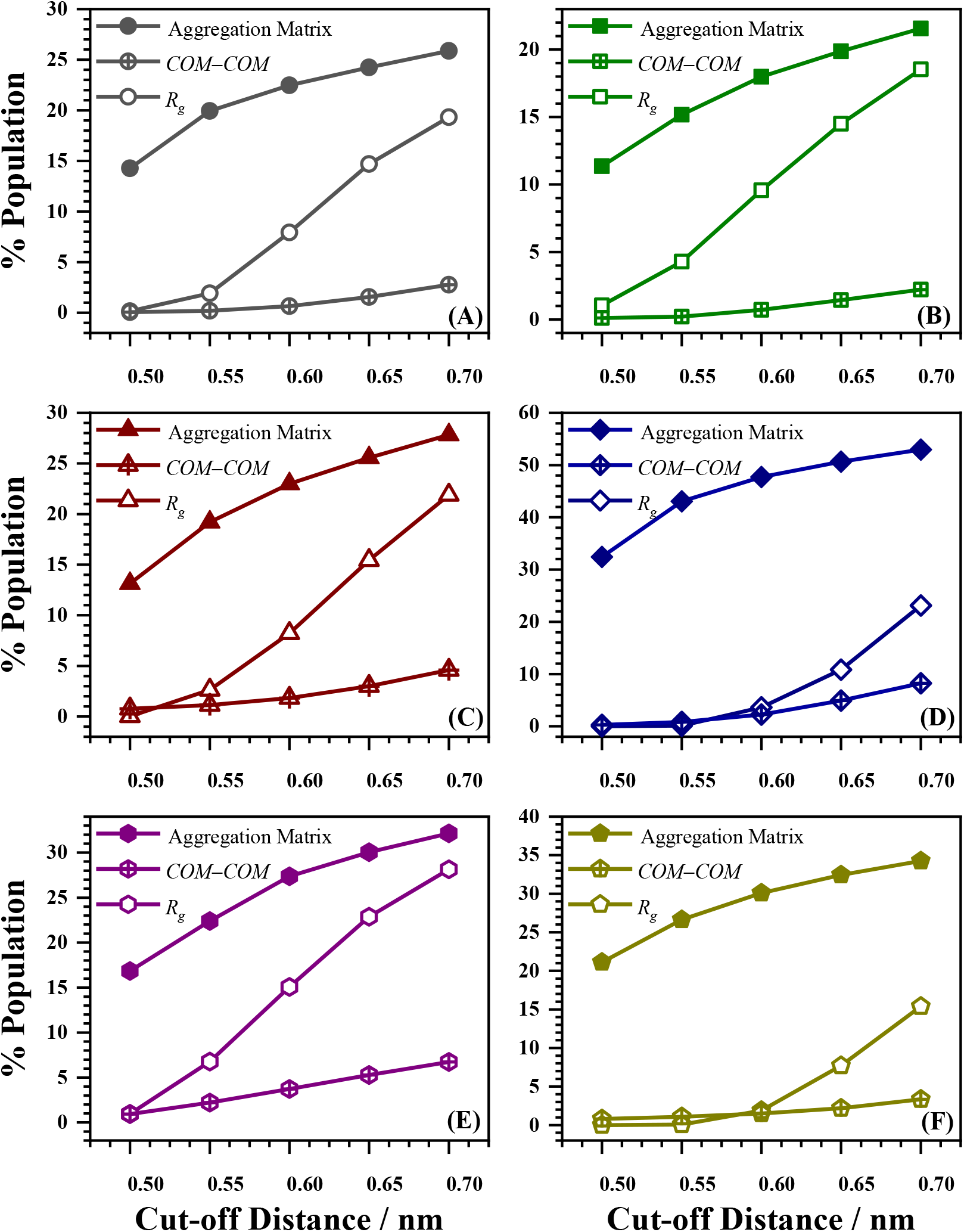
Comparison of dimer formation using the aggregation matrix method (with methods radius of gyration (*R*_*g*_) and center-of-mass to center-of-mass (*COM*–*COM*) methods for (A) NKGAII, (B) STGGYG, (C) SYSGYS, (D) SYSSYGQS, (E) GYNGFG, and (F) GFGNFGTS peptides.

**Figure 8.**
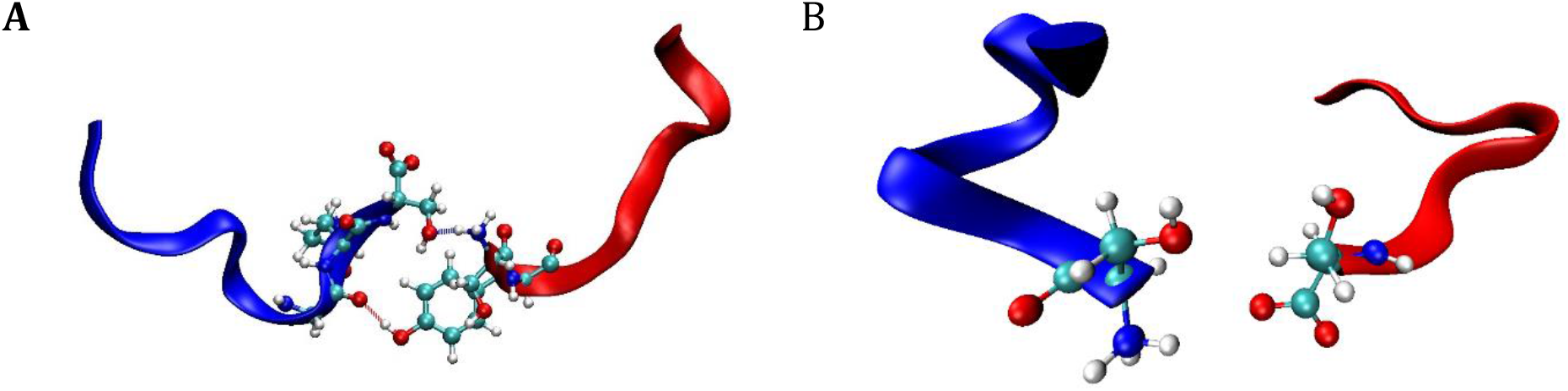
Formation of end-to-end SYSSYGQS dimer captured by the aggregation matrix method. For dimer (A) the values of *R*_*g*_ and *COM*–*COM* are 1.09 and 1.86 nm, respectively, while the corresponding values of the dimer (B) are 0.91 and 1.51 nm, respectively. A *C*_*α*_–*C*_*α*_ cut-off distance of 0.6 nm was used to generate these two structures. These structures might be overlooked depending on the cut-off values of *R*_*g*_ used for analysis. The structure (A) shows the presence of two inter-peptide hydrogen bonds, while structure (B) is a non-specific aggregate.

## CONCLUSIONS

The aggregation propensity of six LCD peptides to form dimers, trimers and tetramers was examined using a newly developed aggregation matrix methodology. In this method, an aggregation matrix is generated by considering the inter-peptide *C*_*α*_–*C*_*α*_ cut-off distances which are encoded to 0 and 1 depending on the distance cut-off. The total dimer population was obtained by counting the number of frames with an aggregation number of at least one. Further, the ordered conformations such as parallel/anti-parallel or shifted parallel/anti-parallel were obtained by analyzing the diagonal/anti-diagonal and shifted diagonal/ani-diagonal, respectively. In a system consisting of three peptides units, the trimerization was analyzed by decomposing the system into three pairs of dimers and the formation of the trimer was deciphered based on snapshots/frames with at least two sets of dimers. Similarly, the tetramer was analyzed by decomposing the system into five dimers and tetramer formation was obtained when two mutually exclusive dimers are within cut-off distances. The explicit comparison of aggregation matrix with the conventional methods such as *COM*–*COM* distance and *R*_*g*_ shows that the conventional methods underestimate the propensity of aggregation. Moreover, the conventional methods do not structurally classify structures such as parallel and anti-parallel, which the aggregation matrix method does by examining matrix elements. The present aggregation matrix is an easy, convenient and inexpensive method to analyze aggregation propensity in the MD trajectory which uses only *nxn* two-dimensional matrices even for systems consisting of several peptide units.

## Supporting Information

Plots showing Hierarchy of aggregation and tables containing population data of ordered and random configurations (PDF).

## Acknowledgments

AT thanks University Grants Commission (UGC) for the research fellowship and RKS thanks IIT Bombay for the institute postdoctoral fellowship. Authors gratefully acknowledge the SpaceTime-2 supercomputing facility at IIT Bombay for the computing time. The support and the resources provided by ‘PARAM Brahma Facility’ under the National Supercomputing Mission, Government of India at the Indian Institute of Science Education and Research (IISER) Pune are gratefully acknowledged. The authors wish to thank Dr. Atanu Das of CSIR-NCL for critical reading and suggestions.

## Notes

### Competing Interest Statement

The authors have declared no competing interest.

### Summary of Updates

We have updated the manuscript by including the analysis of side-chain interactions and comparing them with the Cα aggregation matrices.

