## Supplementary material for "A Binary Matrix Method to Enumerate, Hierarchically Order and Structurally Classify Peptide Aggregation": Supplimentary Figure and Tables

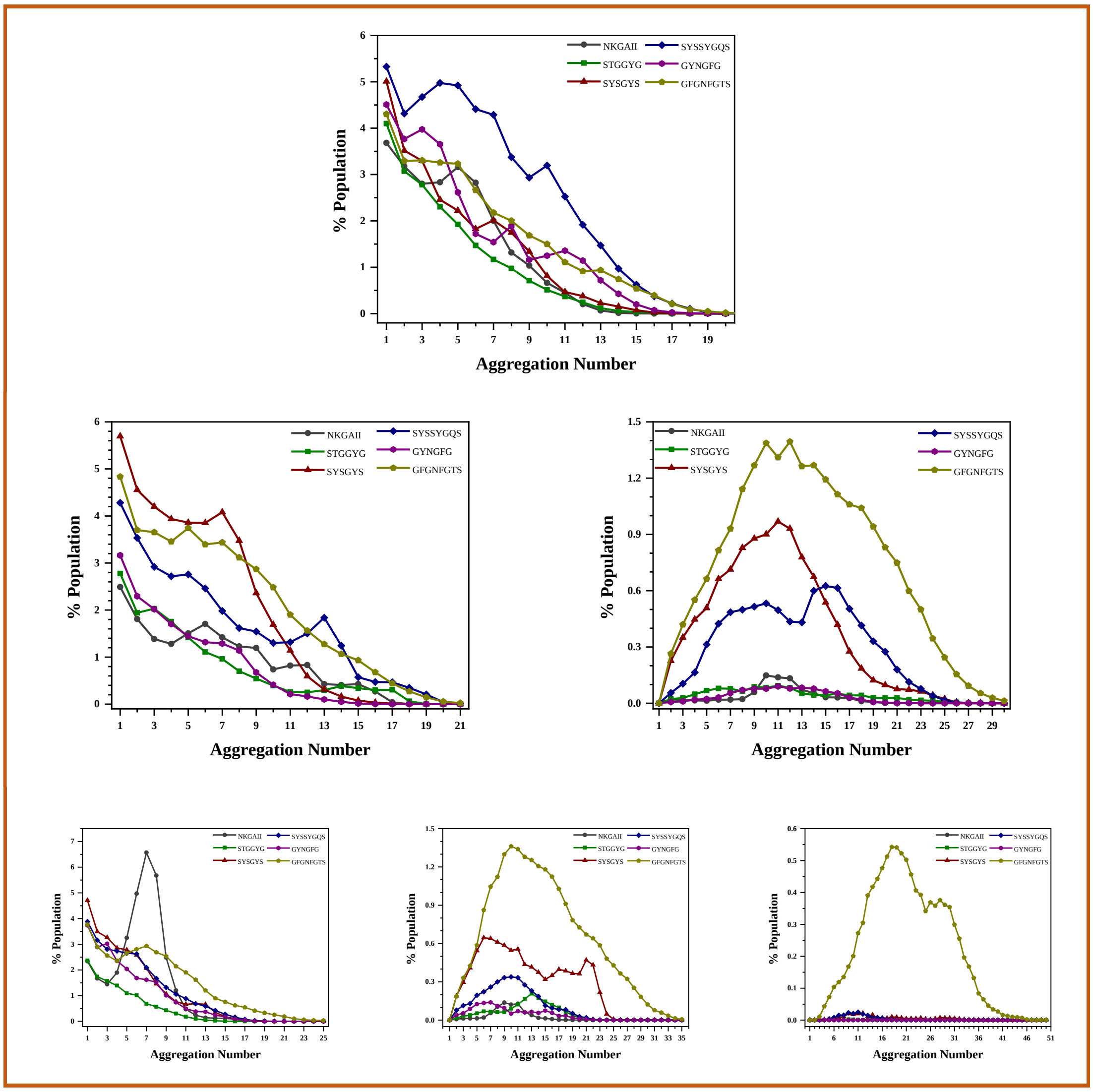


**Figure S1.** (Top panel) Hierarchy of aggregation for the dimer fomation in system containing two peptide units. (Middle panel) Hierarchy of aggregation for the dimer (left) and timer (right) fomation in system containing three peptide units. (Bottom panel) Hierarchy of aggregation for the dimer (left), timer (middle) and tetramer (right) fomation in system containing four peptide units. The $C_\alpha$–$C_\alpha$ cut-off distance of 0.65 nm was used. Notice that for the trimer and the tetramer formation the aggregation number has minimum value of 1 and 2, respectively.

**Table S1: Total population (%) along with the population of ordered and random configurations at various inter-peptide $C_\alpha$–$C_\alpha$ cut-off distances (nm) for two peptide system.**

NKGAII

| Cut-off | Total | Random | Ordered | Parallel | Anti-parallel |
|---|---|---|---|---|---|
| 0.50 | 14.26 | 14.23 | 0.04 | 0.01 | 0.03 |
| 0.55 | 19.94 | 19.45 | 0.49 | 0.11 | 0.39 |
| 0.60 | 22.48 | 20.45 | 2.02 | 0.49 | 1.61 |
| 0.65 | 24.24 | 20.10 | 4.14 | 1.18 | 3.36 |
| 0.70 | 25.86 | 19.72 | 6.14 | 2.03 | 5.02 |
| 0.75 | 27.34 | 18.75 | 8.58 | 3.44 | 7.05 |

STGGYG

| Cut-off | Total | Random | Ordered | Parallel | Anti-parallel |
|---|---|---|---|---|---|
| 0.50 | 11.37 | 11.20 | 0.17 | 0.02 | 0.15 |
| 0.55 | 15.18 | 14.35 | 0.83 | 0.15 | 0.69 |
| 0.60 | 17.99 | 16.14 | 1.84 | 0.54 | 1.41 |
| 0.65 | 19.87 | 16.60 | 3.27 | 1.29 | 2.45 |
| 0.70 | 21.56 | 16.86 | 4.70 | 2.33 | 3.51 |
| 0.75 | 23.06 | 16.53 | 6.54 | 3.81 | 4.92 |

SYSGYS

| Cut-off | Total | Random | Ordered | Parallel | Anti-parallel |
|---|---|---|---|---|---|
| 0.50 | 13.13 | 12.95 | 0.18 | 0.09 | 0.09 |
| 0.55 | 19.21 | 17.14 | 2.07 | 0.80 | 1.28 |
| 0.60 | 23.00 | 18.81 | 4.19 | 1.68 | 2.68 |
| 0.65 | 25.57 | 19.60 | 5.98 | 2.63 | 3.92 |
| 0.70 | 27.82 | 20.03 | 7.79 | 3.80 | 5.14 |
| 0.75 | 29.75 | 19.46 | 10.29 | 5.59 | 6.85 |

SYSSYGQS

| Cut-off | Total | Random | Ordered | Parallel | Anti-parallel |
|---|---|---|---|---|---|
| 0.50 | 32.42 | 31.87 | 0.55 | 0.20 | 0.35 |
| 0.55 | 43.06 | 37.52 | 5.54 | 1.47 | 4.20 |
| 0.60 | 47.73 | 34.85 | 12.88 | 3.59 | 10.43 |
| 0.65 | 50.65 | 32.44 | 18.22 | 6.10 | 14.77 |
| 0.70 | 52.96 | 29.91 | 23.05 | 9.33 | 18.68 |
| 0.75 | 54.69 | 26.60 | 28.09 | 14.75 | 22.91 |

**Table S1 Continued…**

GYNGFG

| Cut-off | Total | Random | Ordered | Parallel | Anti-parallel |
|---|---|---|---|---|---|
| 0.50 | 16.85 | 15.53 | 1.32 | 0.08 | 1.24 |
| 0.55 | 22.40 | 16.20 | 6.21 | 0.39 | 5.89 |
| 0.60 | 27.37 | 19.17 | 8.19 | 0.93 | 7.67 |
| 0.65 | 30.04 | 20.29 | 9.75 | 1.55 | 9.01 |
| 0.70 | 32.15 | 19.67 | 12.48 | 2.65 | 11.49 |
| 0.75 | 33.94 | 18.50 | 15.44 | 4.53 | 14.05 |

NFGGFGTS

| Cut-off | Total | Random | Ordered | Parallel | Anti-parallel |
|---|---|---|---|---|---|
| 0.50 | 21.11 | 20.65 | 0.46 | 0.10 | 0.36 |
| 0.55 | 26.64 | 23.34 | 3.29 | 0.81 | 2.56 |
| 0.60 | 30.10 | 23.17 | 6.93 | 2.20 | 5.45 |
| 0.65 | 32.43 | 22.51 | 9.92 | 4.30 | 7.90 |
| 0.70 | 34.24 | 21.48 | 12.77 | 6.64 | 10.23 |
| 0.75 | 35.82 | 19.72 | 16.09 | 9.81 | 12.94 |

**Tables S2: Dimer population (%) for various peptide consisting of two peptide units in cumulative time segments of 400 ns with cut-off distance of 0.55 nm.**

| **Time (ns)** | NKGAII | STGGYG | SYSGYS | SYSSYGQS | GYNGFG | GFGNFGTS |
|---|---|---|---|---|---|---|
| 0-400 | 22.0 | 10.8 | 26.9 | 34.4 | 29.6 | 21.7 |
| 0-800 | 22.2 | 10.9 | 21.7 | 47.1 | 27.3 | 28.6 |
| 0-1200 | 18.0 | 13.1 | 20.0 | 49.2 | 24.4 | 26.2 |
| 0-1600 | 18.1 | 14.7 | 19.1 | 47.2 | 20.9 | 25.7 |
| 0-2000 | 19.9 | 15.2 | 19.2 | 43.1 | 22.4 | 26.6 |

**Tables S3: Comparison of total population (%) obtained from aggregation matrix with $C_\alpha$–$C_\alpha$ cut-off distance of 0.55 nm and hydrogen bonding.**

| **System** | **Population (Aggregation Matrix)** | **Population (Hydrogen Bonding)** |
|---|---|---|
| NKGAII | 19.9 | 21.4 |
| STGGYG | 15.2 | 12.0 |
| SYSGYS | 19.2 | 21.1 |
| SYSSYGQS | 43.1 | 45.2 |
| GYNGFG | 22.4 | 19.6 |
| NFGGFGTS | 26.6 | 23.5 |

**Tables S4: Comparison of $C_\alpha$ aggregation matrix (left) with the corresponding HB matrix (right) for the structures shown in Figure 2.**

**Parallel**

| **Cα** | B1 | B2 | B3 | B4 | B5 | B6 | B7 | B8 | **HB** | B1 | B2 | B3 | B4 | B5 | B6 | B7 | B8 |
|---|---|---|---|---|---|---|---|---|---|---|---|---|---|---|---|---|---|
| A1 | **0** | 0 | 0 | 0 | 0 | 0 | 0 | 0 | A1 | 0 | 0 | 0 | 0 | 0 | 0 | 0 | 0 |
| A2 | 0 | **1** | 1 | 0 | 0 | 0 | 0 | 0 | A2 | 0 | 0 | 0 | 0 | 0 | 0 | 0 | 0 |
| A3 | 0 | 0 | **1** | 0 | 0 | 0 | 0 | 0 | A3 | 0 | **1** | 0 | **1** | 0 | 0 | 0 | 0 |
| A4 | 0 | 0 | 0 | **1** | 1 | 0 | 0 | 0 | A4 | 0 | 0 | 0 | 0 | 0 | 0 | 0 | 0 |
| A5 | 0 | 0 | 0 | 0 | **1** | 0 | 0 | 0 | A5 | 0 | 0 | 0 | 0 | 0 | 0 | 0 | 0 |
| A6 | 0 | 0 | 0 | 0 | 0 | **0** | 0 | 0 | A6 | 0 | 0 | 0 | 0 | 0 | 0 | 0 | 0 |
| A7 | 0 | 0 | 0 | 0 | 0 | 0 | **0** | 0 | A7 | 0 | 0 | 0 | 0 | 0 | **1** | 0 | 0 |
| A8 | 0 | 0 | 0 | 0 | 0 | 0 | 0 | **0** | A8 | 0 | 0 | 0 | 0 | 0 | 0 | 0 | 0 |

**Anti-parallel**

| **Cα** | B1 | B2 | B3 | B4 | B5 | B6 | B7 | B8 | **HB** | B1 | B2 | B3 | B4 | B5 | B6 | B7 | B8 |
|---|---|---|---|---|---|---|---|---|---|---|---|---|---|---|---|---|---|
| A1 | 0 | 0 | 0 | 0 | 0 | 0 | 1 | **1** | A1 | 0 | 0 | 0 | 0 | 0 | 0 | 0 | **1** |
| A2 | 0 | 0 | 0 | 0 | 0 | 0 | **1** | 1 | A2 | 0 | 0 | 0 | 0 | 0 | 0 | 0 | 0 |
| A3 | 0 | 0 | 0 | 0 | 0 | **1** | 1 | 1 | A3 | 0 | 0 | 0 | **1** | 0 | 0 | 0 | 0 |
| A4 | 0 | 0 | 0 | 0 | **0** | 0 | 0 | 0 | A4 | 0 | 0 | 0 | 0 | 0 | 0 | 0 | **1** |
| A5 | 0 | 0 | 1 | **1** | 0 | 0 | 0 | 0 | A5 | 0 | 0 | 0 | 0 | 0 | 0 | 0 | 0 |
| A6 | 0 | 1 | **1** | 1 | 0 | 0 | 0 | 0 | A6 | 0 | 0 | **1** | 0 | 0 | 0 | 0 | 0 |
| A7 | 1 | **1** | 0 | 0 | 0 | 0 | 0 | 0 | A7 | 0 | **1** | 0 | 0 | 0 | 0 | 0 | 0 |
| A8 | **1** | 0 | 0 | 0 | 0 | 0 | 0 | 0 | A8 | 0 | 0 | 0 | 0 | 0 | 0 | 0 | 0 |

**Shifted parallel (down)**

| **Cα** | B1 | B2 | B3 | B4 | B5 | B6 | B7 | B8 | **HB** | B1 | B2 | B3 | B4 | B5 | B6 | B7 | B8 |
|---|---|---|---|---|---|---|---|---|---|---|---|---|---|---|---|---|---|
| A1 | 0 | 0 | 0 | 0 | 0 | 0 | 0 | 0 | A1 | 0 | 0 | 0 | 0 | 0 | 0 | 0 | 0 |
| A2 | **1** | 1 | 0 | 0 | 0 | 0 | 0 | 0 | A2 | **1** | 0 | 0 | 0 | 0 | 0 | 0 | 0 |
| A3 | 1 | **1** | 0 | 0 | 0 | 0 | 0 | 0 | A3 | 0 | 0 | 0 | 0 | 0 | 0 | 0 | 0 |
| A4 | 0 | 1 | **1** | 0 | 0 | 0 | 0 | 0 | A4 | 0 | 0 | 1 | 0 | 0 | 0 | 0 | 0 |
| A5 | 0 | 0 | 0 | **0** | 0 | 0 | 0 | 0 | A5 | 0 | 0 | 0 | 0 | 0 | 0 | 0 | 0 |
| A6 | 0 | 0 | 0 | 0 | **0** | 1 | 0 | 0 | A6 | 0 | 0 | 0 | 0 | 0 | 0 | 0 | 0 |
| A7 | 0 | 0 | 0 | 0 | 0 | **1** | 0 | 0 | A7 | 0 | **1** | **1** | 0 | 0 | 0 | 0 | 0 |
| A8 | 0 | 0 | 0 | 0 | 0 | 0 | **0** | 0 | A8 | 0 | 0 | 0 | 0 | 0 | 0 | 0 | 0 |

**Shifted anti-parallel (down)**

| **Cα** | B1 | B2 | B3 | B4 | B5 | B6 | B7 | B8 | **HB** | B1 | B2 | B3 | B4 | B5 | B6 | B7 | B8 |
|---|---|---|---|---|---|---|---|---|---|---|---|---|---|---|---|---|---|
| A1 | 0 | 0 | 0 | 0 | 0 | 0 | 0 | 0 | A1 | 0 | 0 | 0 | 0 | 0 | 0 | 0 | 0 |
| A2 | 0 | 0 | 0 | 0 | 0 | 0 | 0 | **1** | A2 | 0 | 0 | 0 | 0 | 0 | 0 | 0 | 0 |
| A3 | 0 | 0 | 0 | 0 | 0 | 1 | **1** | 1 | A3 | 0 | 0 | 0 | **1** | 0 | 0 | 0 | 0 |
| A4 | 0 | 0 | 0 | 0 | 1 | **1** | 1 | 0 | A4 | 0 | 0 | 0 | 0 | 0 | 0 | 0 | 0 |
| A5 | 0 | 0 | 0 | 0 | **1** | 0 | 0 | 0 | A5 | 0 | 0 | **1** | 0 | 0 | 0 | 0 | 0 |
| A6 | 0 | 0 | 1 | **1** | 1 | 0 | 0 | 0 | A6 | 0 | 0 | 0 | 0 | 0 | 0 | 0 | 0 |
| A7 | 0 | 0 | **1** | 0 | 0 | 0 | 0 | 0 | A7 | 0 | 0 | 0 | 0 | **1** | 0 | 0 | 0 |
| A8 | 0 | **1** | 1 | 0 | 0 | 0 | 0 | 0 | A8 | 0 | 0 | 0 | 0 | 0 | 0 | 0 | 0 |

**Shifted parallel (up)**

| **Cα** | B1 | B2 | B3 | B4 | B5 | B6 | B7 | B8 | **HB** | B1 | B2 | B3 | B4 | B5 | B6 | B7 | B8 |
|---|---|---|---|---|---|---|---|---|---|---|---|---|---|---|---|---|---|
| A1 | 0 | **1** | 0 | 0 | 0 | 0 | 0 | 0 | A1 | 0 | 0 | 0 | 0 | 0 | 0 | 0 | 0 |
| A2 | 0 | 1 | **1** | 0 | 0 | 0 | 0 | 0 | A2 | 0 | 0 | 0 | 0 | 0 | 0 | 0 | **1** |
| A3 | 0 | 0 | 1 | **1** | 0 | 0 | 0 | 0 | A3 | 0 | 0 | 0 | 0 | 0 | 0 | 0 | 0 |
| A4 | 0 | 0 | 1 | 1 | **1** | 1 | 0 | 0 | A4 | 0 | 0 | **1** | 0 | 0 | **1** | **1** | 0 |
| A5 | 0 | 0 | 0 | 0 | 0 | **0** | 0 | 0 | A5 | 0 | 0 | 0 | 0 | 0 | 0 | 0 | 0 |
| A6 | 0 | 0 | 0 | 0 | 0 | 0 | **0** | 0 | A6 | 0 | 0 | 0 | 0 | 0 | 0 | 0 | 0 |
| A7 | 0 | 0 | 0 | 0 | 0 | 0 | 0 | **0** | A7 | 0 | 0 | 0 | 0 | 0 | 0 | 0 | 0 |
| A8 | 0 | 0 | 0 | 0 | 0 | 0 | 0 | 1 | A8 | 0 | 0 | 0 | 0 | 0 | 0 | 0 | 0 |

**Shifted anti-parallel (up)**

| Cα | $B_1$ | $B_2$ | $B_3$ | $B_4$ | $B_5$ | $B_6$ | $B_7$ | $B_8$ | HB | $B_1$ | $B_2$ | $B_3$ | $B_4$ | $B_5$ | $B_6$ | $B_7$ | $B_8$ |
|---|---|---|---|---|---|---|---|---|---|---|---|---|---|---|---|---|---|
| $A_1$ | 0 | 0 | 0 | 0 | 1 | 1 | **0** | 0 | $A_1$ | 0 | 0 | 0 | 0 | **1** | 0 | 0 | 0 |
| $A_2$ | 0 | 0 | 0 | 0 | 1 | **1** | 0 | 0 | $A_2$ | 0 | 0 | 0 | 0 | 0 | 0 | 0 | 0 |
| $A_3$ | 0 | 0 | 0 | 0 | **1** | 1 | 0 | 0 | $A_3$ | 0 | 0 | **1** | 0 | 0 | 0 | 0 | 0 |
| $A_4$ | 0 | 0 | 0 | **0** | 0 | 0 | 0 | 0 | $A_4$ | 0 | 0 | 0 | 0 | 0 | 0 | 0 | 0 |
| $A_5$ | 0 | 0 | **1** | 0 | 0 | 0 | 0 | 0 | $A_5$ | 0 | 0 | 0 | 0 | 0 | 0 | 0 | 0 |
| $A_6$ | 1 | **1** | 1 | 0 | 0 | 0 | 0 | 0 | $A_6$ | **1** | 0 | 0 | 0 | 0 | 0 | 0 | 0 |
| $A_7$ | **1** | 1 | 1 | 0 | 0 | 0 | 0 | 0 | $A_7$ | 0 | 0 | 0 | 0 | 0 | 0 | 0 | 0 |
| $A_8$ | 1 | 0 | 0 | 0 | 0 | 0 | 0 | 0 | $A_8$ | **1** | 0 | 0 | 0 | 0 | 0 | 0 | 0 |

**T-shaped**

| Cα | $B_1$ | $B_2$ | $B_3$ | $B_4$ | $B_5$ | $B_6$ | $B_7$ | $B_8$ | HB | $B_1$ | $B_2$ | $B_3$ | $B_4$ | $B_5$ | $B_6$ | $B_7$ | $B_8$ |
|---|---|---|---|---|---|---|---|---|---|---|---|---|---|---|---|---|---|
| $A_1$ | 0 | 0 | 0 | 0 | 0 | 0 | 0 | 0 | $A_1$ | 0 | 0 | 0 | 0 | 0 | 0 | 0 | 0 |
| $A_2$ | 0 | 0 | 0 | 0 | 0 | 0 | 0 | 0 | $A_2$ | 0 | 0 | 0 | 0 | 0 | 0 | 0 | 0 |
| $A_3$ | 0 | 0 | 0 | 0 | 0 | 0 | 0 | 0 | $A_3$ | 0 | 0 | 0 | 0 | 0 | 0 | 0 | 0 |
| $A_4$ | 0 | 0 | 0 | 0 | 0 | 0 | 0 | 0 | $A_4$ | 0 | 0 | 0 | 0 | 0 | 0 | 0 | 0 |
| $A_5$ | **1** | 0 | 0 | 0 | 0 | 0 | 0 | 0 | $A_5$ | **1** | 0 | 0 | 0 | 0 | 0 | 0 | 0 |
| $A_6$ | 0 | 0 | 0 | 0 | 0 | 0 | 0 | 0 | $A_6$ | 0 | 0 | 0 | 0 | 0 | 0 | 0 | 0 |
| $A_7$ | 0 | 0 | 0 | 0 | 0 | 0 | 0 | 0 | $A_7$ | 0 | 0 | 0 | 0 | 0 | 0 | 0 | 0 |
| $A_8$ | 0 | 0 | 0 | 0 | 0 | 0 | 0 | 0 | $A_8$ | 0 | 0 | 0 | 0 | 0 | 0 | 0 | 0 |

**X-shaped**

| Cα | $B_1$ | $B_2$ | $B_3$ | $B_4$ | $B_5$ | $B_6$ | $B_7$ | $B_8$ | HB | $B_1$ | $B_2$ | $B_3$ | $B_4$ | $B_5$ | $B_6$ | $B_7$ | $B_8$ |
|---|---|---|---|---|---|---|---|---|---|---|---|---|---|---|---|---|---|
| $A_1$ | 0 | 0 | 0 | 0 | 0 | 0 | 0 | 0 | $A_1$ | 0 | 0 | 0 | 0 | 0 | 0 | 0 | 0 |
| $A_2$ | 0 | 0 | 0 | 0 | 0 | 0 | 0 | 0 | $A_2$ | 0 | **1** | 0 | 0 | 0 | 0 | 0 | 0 |
| $A_3$ | 0 | 0 | 0 | 0 | 0 | 0 | 0 | 0 | $A_3$ | 0 | 0 | 0 | 0 | 0 | 0 | 0 | 0 |
| $A_4$ | 0 | 0 | 0 | **1** | **1** | 0 | 0 | 0 | $A_4$ | 0 | 0 | 0 | 0 | 0 | 0 | 0 | 0 |
| $A_5$ | 0 | 0 | 0 | **1** | **1** | 0 | 0 | 0 | $A_5$ | 0 | 0 | 0 | 0 | 0 | 0 | 0 | 0 |
| $A_6$ | 0 | 0 | 0 | 0 | 0 | 0 | 0 | 0 | $A_6$ | 0 | 0 | 0 | 0 | 0 | 0 | 0 | 0 |
| $A_7$ | 0 | 0 | 0 | 0 | 0 | 0 | 0 | 0 | $A_7$ | 0 | 0 | 0 | 0 | 0 | 0 | 0 | 0 |
| $A_8$ | 0 | 0 | 0 | 0 | 0 | 0 | 0 | 0 | $A_8$ | 0 | 0 | 0 | 0 | 0 | 0 | 0 | 0 |

**Tables S5: Total population (%) along with the population of ordered and random configurations at various inter-peptide $C_\alpha$–$C_\alpha$ cut-off distances (nm) for three peptide system.**

NKGAII

| Cut-off | Dimer | | | Trimer | | |
|---|---|---|---|---|---|---|
| | Total | Random | Order | Total | Random | Order |
| 0.50 | 12.7 | 12.3 | 0.3 | 0.2 | 0.2 | 0.0 |
| 0.55 | 15.5 | 12.6 | 2.9 | 0.6 | 0.6 | 0.0 |
| 0.60 | 17.0 | 12.2 | 4.7 | 0.7 | 0.7 | 0.0 |
| 0.65 | 18.0 | 14.1 | 3.9 | 0.8 | 0.8 | 0.0 |
| 0.70 | 18.9 | 14.5 | 4.4 | 0.9 | 0.9 | 0.1 |
| 0.75 | 19.8 | 14.6 | 5.2 | 1.0 | 0.9 | 0.1 |

**Table S5 Continued…**

STGGYG

| Cut-off | Dimer | | | Trimer | | |
|---|---|---|---|---|---|---|
| | Total | Random | Order | Total | Random | Order |
| 0.50 | 10.3 | 9.2 | 1.1 | 0.5 | 0.5 | 0.0 |
| 0.55 | 12.7 | 9.8 | 2.9 | 0.8 | 0.8 | 0.0 |
| 0.60 | 14.5 | 10.9 | 3.5 | 1.0 | 1.0 | 0.0 |
| 0.65 | 15.8 | 12.0 | 3.9 | 1.2 | 1.1 | 0.0 |
| 0.70 | 16.9 | 12.8 | 4.2 | 1.3 | 1.2 | 0.1 |
| 0.75 | 17.9 | 13.6 | 4.3 | 1.4 | 1.3 | 0.1 |

SYSGYS

| Cut-off | Dimer | | | Trimer | | |
|---|---|---|---|---|---|---|
| | Total | Random | Order | Total | Random | Order |
| 0.50 | 28.5 | 27.7 | 0.8 | 4.2 | 4.2 | 0.0 |
| 0.55 | 35.6 | 31.2 | 4.4 | 7.5 | 7.4 | 0.1 |
| 0.60 | 38.4 | 30.3 | 8.0 | 9.5 | 9.1 | 0.4 |
| 0.65 | 40.0 | 28.6 | 11.5 | 10.8 | 9.4 | 1.4 |
| 0.70 | 41.4 | 26.8 | 14.6 | 12.1 | 9.0 | 3.1 |
| 0.75 | 42.6 | 25.3 | 17.2 | 13.3 | 7.6 | 5.7 |

SYSSYGQS

| Cut-off | Dimer | | | Trimer | | |
|---|---|---|---|---|---|---|
| | Total | Random | Order | Total | Random | Order |
| 0.50 | 24.7 | 22.5 | 2.2 | 3.9 | 3.9 | 0.0 |
| 0.55 | 29.0 | 23.2 | 5.8 | 6.3 | 6.3 | 0.0 |
| 0.60 | 31.5 | 24.2 | 7.4 | 7.5 | 7.5 | 0.1 |
| 0.65 | 33.2 | 24.4 | 8.8 | 8.2 | 8.1 | 0.2 |
| 0.70 | 34.4 | 24.5 | 9.9 | 8.8 | 8.5 | 0.4 |
| 0.75 | 35.5 | 25.6 | 9.9 | 9.2 | 8.2 | 1.0 |

GYNGFG

| Cut-off | Dimer | | | Trimer | | |
|---|---|---|---|---|---|---|
| | Total | Random | Order | Total | Random | Order |
| 0.50 | 9.6 | 9.3 | 0.3 | 0.5 | 0.5 | 0.0 |
| 0.55 | 12.3 | 11.0 | 1.3 | 0.7 | 0.7 | 0.0 |
| 0.60 | 14.4 | 12.1 | 2.3 | 0.8 | 0.8 | 0.0 |
| 0.65 | 16.0 | 12.7 | 3.3 | 0.9 | 0.9 | 0.0 |
| 0.70 | 17.5 | 13.1 | 4.5 | 0.9 | 0.9 | 0.0 |
| 0.75 | 18.8 | 13.6 | 5.3 | 1.0 | 0.9 | 0.1 |

**Table S5 Continued…**

NFGGFGTS

| Cut-off | Dimer | | | Trimer | | |
|---|---|---|---|---|---|---|
| | Total | Random | Order | Total | Random | Order |
| 0.50 | 39.6 | 38.1 | 1.5 | 10.6 | 10.6 | 0.0 |
| 0.55 | 42.1 | 33.6 | 8.5 | 15.5 | 15.4 | 0.1 |
| 0.60 | 42.7 | 28.8 | 13.9 | 19.2 | 18.6 | 0.6 |
| 0.65 | 43.0 | 26.1 | 16.9 | 21.6 | 20.3 | 1.3 |
| 0.70 | 43.2 | 24.7 | 18.5 | 23.7 | 21.3 | 2.4 |
| 0.75 | 43.3 | 25.8 | 17.5 | 25.3 | 20.9 | 4.5 |

**Tables S6: Total population (%) along with the population of ordered and random configurations at various inter-peptide $C_\alpha$–$C_\alpha$ cut-off distances (nm) for four peptide system.**

NKGAII

| Cut-off | Dimer | | | Trimer | | | Tetramer | | |
|---|---|---|---|---|---|---|---|---|---|
| | Total | Random | Order | Total | Random | Order | Total | Random | Order |
| 0.50 | 23.3 | 23.1 | 0.2 | 0.2 | 0.0 | 0.0 | 0 | 0 | 0 |
| 0.55 | 30.6 | 29.5 | 1.1 | 0.5 | 0.0 | 0.0 | 0 | 0 | 0 |
| 0.60 | 31.9 | 29.2 | 2.7 | 0.7 | 0.0 | 0.0 | 0 | 0 | 0 |
| 0.65 | 32.7 | 27.7 | 5.0 | 0.8 | 0.0 | 0.0 | 0 | 0 | 0 |
| 0.70 | 33.5 | 23.3 | 10.2 | 0.9 | 0.0 | 0.0 | 0 | 0 | 0 |
| 0.75 | 34.2 | 16.9 | 17.3 | 1.0 | 0.0 | 0.0 | 0 | 0 | 0 |

STGGYG

| Cut-off | Dimer | | | Trimer | | | Tetramer | | |
|---|---|---|---|---|---|---|---|---|---|
| | Total | Random | Order | Total | Random | Order | Total | Random | Order |
| 0.50 | 7.2 | 7.1 | 0.1 | 1.1 | 1.1 | 0.0 | 0 | 0 | 0 |
| 0.55 | 9.0 | 8.5 | 0.4 | 1.1 | 1.1 | 0.0 | 0 | 0 | 0 |
| 0.60 | 10.5 | 9.2 | 1.3 | 1.3 | 1.3 | 0.0 | 0 | 0 | 0 |
| 0.65 | 11.5 | 9.1 | 2.4 | 1.3 | 1.3 | 0.0 | 0 | 0 | 0 |
| 0.70 | 12.4 | 8.9 | 3.5 | 1.5 | 1.5 | 0.0 | 0 | 0 | 0 |
| 0.75 | 13.2 | 8.6 | 4.6 | 1.5 | 1.5 | 0.0 | 0 | 0 | 0 |

**Table S6 Continued…**

SYSGYS

| Cut-off | Dimer | | | Trimer | | | Tetramer | | |
|---|---|---|---|---|---|---|---|---|---|
| | Total | Random | Order | Total | Random | Order | Total | Random | Order |
| 0.50 | 19.8 | 19.4 | 0.4 | 4.8 | 4.8 | 0.0 | 0.1 | 0 | 0 |
| 0.55 | 23.9 | 20.0 | 3.8 | 7.3 | 7.3 | 0.0 | 0.2 | 0 | 0 |
| 0.60 | 26.1 | 18.3 | 7.8 | 8.7 | 8.7 | 0.1 | 0.2 | 0 | 0 |
| 0.65 | 27.8 | 16.0 | 11.8 | 9.6 | 9.4 | 0.2 | 0.2 | 0 | 0 |
| 0.70 | 29.1 | 13.5 | 15.6 | 10.4 | 10.0 | 0.4 | 0.3 | 0 | 0 |
| 0.75 | 30.3 | 11.9 | 18.4 | 11.0 | 10.3 | 0.7 | 0.3 | 0 | 0 |

SYSSYGQS

| Cut-off | Dimer | | | Trimer | | | Tetramer | | |
|---|---|---|---|---|---|---|---|---|---|
| | Total | Random | Order | Total | Random | Order | Total | Random | Order |
| 0.50 | 17.6 | 17.4 | 0.2 | 0.6 | 0.6 | 0.0 | 0.0 | 0 | 0 |
| 0.55 | 22.5 | 20.7 | 1.8 | 2.3 | 2.3 | 0.0 | 0.1 | 0 | 0 |
| 0.60 | 25.3 | 20.5 | 4.8 | 3.1 | 3.1 | 0.0 | 0.1 | 0 | 0 |
| 0.65 | 27.1 | 19.0 | 8.1 | 3.5 | 3.4 | 0.0 | 0.2 | 0 | 0 |
| 0.70 | 28.5 | 17.1 | 11.4 | 3.8 | 3.7 | 0.1 | 0.2 | 0 | 0 |
| 0.75 | 29.6 | 14.5 | 15.1 | 4.0 | 3.7 | 0.2 | 0.2 | 0 | 0 |

GYNGFG

| Cut-off | Dimer | | | Trimer | | | Tetramer | | |
|---|---|---|---|---|---|---|---|---|---|
| | Total | Random | Order | Total | Random | Order | Total | Random | Order |
| 0.50 | 13.9 | 12.9 | 1.1 | 0.4 | 0.4 | 0.0 | 0.0 | 0 | 0 |
| 0.55 | 17.7 | 15.0 | 2.7 | 0.7 | 0.7 | 0.0 | 0.0 | 0 | 0 |
| 0.60 | 20.5 | 16.3 | 4.2 | 1.0 | 1.0 | 0.0 | 0.0 | 0 | 0 |
| 0.65 | 22.4 | 16.7 | 5.7 | 1.3 | 1.3 | 0.0 | 0.0 | 0 | 0 |
| 0.70 | 24.0 | 16.6 | 7.4 | 1.6 | 1.6 | 0.0 | 0.0 | 0 | 0 |
| 0.75 | 25.1 | 15.6 | 9.5 | 1.9 | 1.8 | 0.1 | 0.0 | 0 | 0 |

GFGNFGTS

| Cut-off | Dimer | | | Trimer | | | Tetramer | | |
|---|---|---|---|---|---|---|---|---|---|
| | Total | Random | Order | Total | Random | Order | Total | Random | Order |
| 0.50 | 37.4 | 34.7 | 2.7 | 15.9 | 15.9 | 0.0 | 4.5 | 4.5 | 0.0 |
| 0.55 | 37.1 | 24.3 | 12.9 | 19.6 | 19.3 | 0.3 | 7.4 | 7.4 | 0.0 |
| 0.60 | 36.5 | 14.0 | 22.5 | 21.3 | 19.6 | 1.7 | 9.4 | 9.3 | 0.1 |
| 0.65 | 36.4 | 7.5 | 28.9 | 22.3 | 18.8 | 3.5 | 10.6 | 10.2 | 0.4 |
| 0.70 | 36.4 | 3.9 | 32.5 | 23.0 | 17.2 | 5.8 | 11.4 | 10.6 | 0.8 |
| 0.75 | 36.4 | 1.2 | 35.2 | 23.4 | 14.9 | 8.5 | 12.2 | 10.6 | 1.6 |
